# Tissue macrophages stem sepsis induced altered organ repair

**DOI:** 10.1101/2025.11.18.689163

**Authors:** Charles de Roquetaillade, Manon Durand, Victor Beaucote, Antoine Borouchaki, Pierre-Louis Blot, Jeremy Guillemin, Sehmi Mansouri, Jovana Broćić, Emeline Cherchame, Marie Coutelier, Meryem Memmadi, Aura Zelco, Malha Sadoune, Adrien Picod, Christos E Chadjichristos, Jane-Lise Samuel, Romane Lafontaine, Nahid Tabibzadeh, Etienne Gayat, Alexandre Mebazaa, Benjamin Glenn Chousterman

## Abstract

**Introduction:** Sepsis, the most severe manifestation of infection, remains both common and life-threatening. Beyond the acute phase, growing evidence highlights that sepsis survivors face an increased risk of chronic organ dysfunction, including kidney and heart failure. While early immune mechanisms of sepsis have been extensively studied, the biological processes underlying delayed post-sepsis organ sequelae remain poorly understood.

**Methods:** Using murine models of sepsis, we investigated the long-term impact of sepsis on organ immune landscapes and tissue repair. Fate-mapping experiments were conducted to trace macrophage origin and persistence. Single-cell RNA sequencing of the heart and kidney was performed to characterize macrophage subpopulations and transcriptional reprogramming following sepsis. Functional consequences of this immune remodeling were assessed by challenging post-septic animals with angiotensin II to evaluate secondary injury susceptibility and fibrotic remodeling.

**Results:** Sepsis induced transient multiorgan failure but led to a sustained expansion of tissue macrophages. Fate-mapping demonstrated that this expansion was largely driven by the recruitment and engraftment of monocyte-derived macrophages. Single-cell transcriptomic analyses revealed that post-septic macrophages acquired a distinct proinflammatory and profibrotic signature, consistent with persistent microenvironmental activation. Functionally, this altered macrophage landscape impaired organ resilience, as evidenced by increased mortality and exacerbated cardiac and renal fibrosis upon secondary challenge.

**Discussion:** Our findings identify macrophage reprogramming as a central mechanism linking sepsis to long-term organ vulnerability. The persistence of monocyte-derived macrophages with maladaptive transcriptional profiles promotes fibrotic remodeling and impaired tissue repair. Targeting macrophage recruitment or reprogramming may represent a promising strategy to prevent chronic organ failure among sepsis survivors.

**SIGNIFICANCE STATEMENT:** Sepsis survivors often develop chronic organ dysfunction, yet the mechanisms linking acute infection to long-term damage remain unclear. We show that sepsis triggers expansion of inflammatory and profibrotic macrophages derived from bone marrow precursors in the kidney and heart. These alterations were associated with impaired organ resilience to subsequent injury, which suggest macrophage-driven maladaptive repair as a potential target to improve long-term outcomes after sepsis.

## INTRODUCTION

Sepsis is defined as a life-threatening organ failure due to a dysregulated host response to an infection (1). It affects annually approximately 45 million individuals with an increasing incidence and is responsible for approximately 11 million deaths annually, which makes it the 3^rd^ most frequent cause of death after cardiovascular diseases and cancer (2). During sepsis, cardiac and renal failure are frequent. Due to improvement in patient management, most patients presenting with sepsis survive the acute episode and may leave hospitals with apparent recovery. However, sepsis has a high mid- and long-term morbidity and mortality (3–5).

Despite apparent recovery, recent studies highlighted that sepsis survivors had impairment in their quality of life and increased risk of developing chronic organ failure and especially cardio-renal diseases (CRD), *i.e.,* chronic kidney disease and heart failure. This accelerated CRD increases the risk of rehospitalization, need for renal replacement therapy (RRT) and increases the risk of death (3–5). These phenomena could be attributable to prolonged systemic immune disturbances.

A hallmark of sepsis and septic shock is massive recruitment of monocytes from various sources including bone marrow and hematopoietic organs, in a process called emergency hematopoiesis (6). Mobilized quickly upon infection, monocytes can differentiate to patrolling monocyte or give rise to monocyte-derived macrophages (7).

Tissue-resident macrophages that multiply independently of emergency hematopoiesis are usually recognized as less inflammatory than monocyte-derived macrophages (8) and originate from other source than monocytes (9). However, depending on the organ and challenge, several authors have observed various phenotypes (8,10). The observed increase in macrophage population after sepsis growth may rely on both monocyte recruitment and local proliferation, although no proper demonstration of this was ever made (11,12). On another level, the global inflammation due to massive cytokine release, together with the high level of circulating pathogen-associated molecular patterns and damage-associated molecular patterns, most likely affect resident macrophages and monocytes in their number and phenotypes. Those two coexisting phenomena most likely have long-term consequences, either bolstering or compromising the organ resilience to subsequent challenges.

In the present study, we focused on the kidney and heart, the two most important organs contributing to post-sepsis outcome and risk of re-hospitalization. Our results show that sepsis causes transient organ failure and induces an increase of tissue macrophages, which is mainly due to recruited monocyte-derived macrophages based on fate mapping. scRNAseq analyses of heart and kidney show that monocyte-derived macrophages have a distinct transcriptomic signature with pro-inflammatory and pro-fibrotic features. This change in tissue macrophages landscape is associated with an increase in mortality and tissue fibrosis when septic mice are undergoing an angiotensin-2 challenge. Our findings suggest that macrophages stem sepsis-induced altered organ repair and could represent a therapeutic target in order to ameliorate septic patient long-term outcomes.

## RESULTS

### Sepsis Induces Transient Multiorgan Dysfunction Without Long-term Fibrosis

To better characterize the long-term consequences of sepsis, we used a sublethal cecal ligation and puncture (CLP) model in 6–10-week-old C57BL/6 mice. This model induced clear clinical signs of sepsis while maintaining a low mortality rate (**Figure 1A**) (13). Mice exhibited significant weight loss (10–15%) and behavioral alterations during the first few days after CLP (clinical scoring details are provided in the **Supplementary Material**). Surviving animals progressively regained normal weight and behavior by day 14. Cumulative mortality reached 20–25% by day 21.

**Figure 1.**
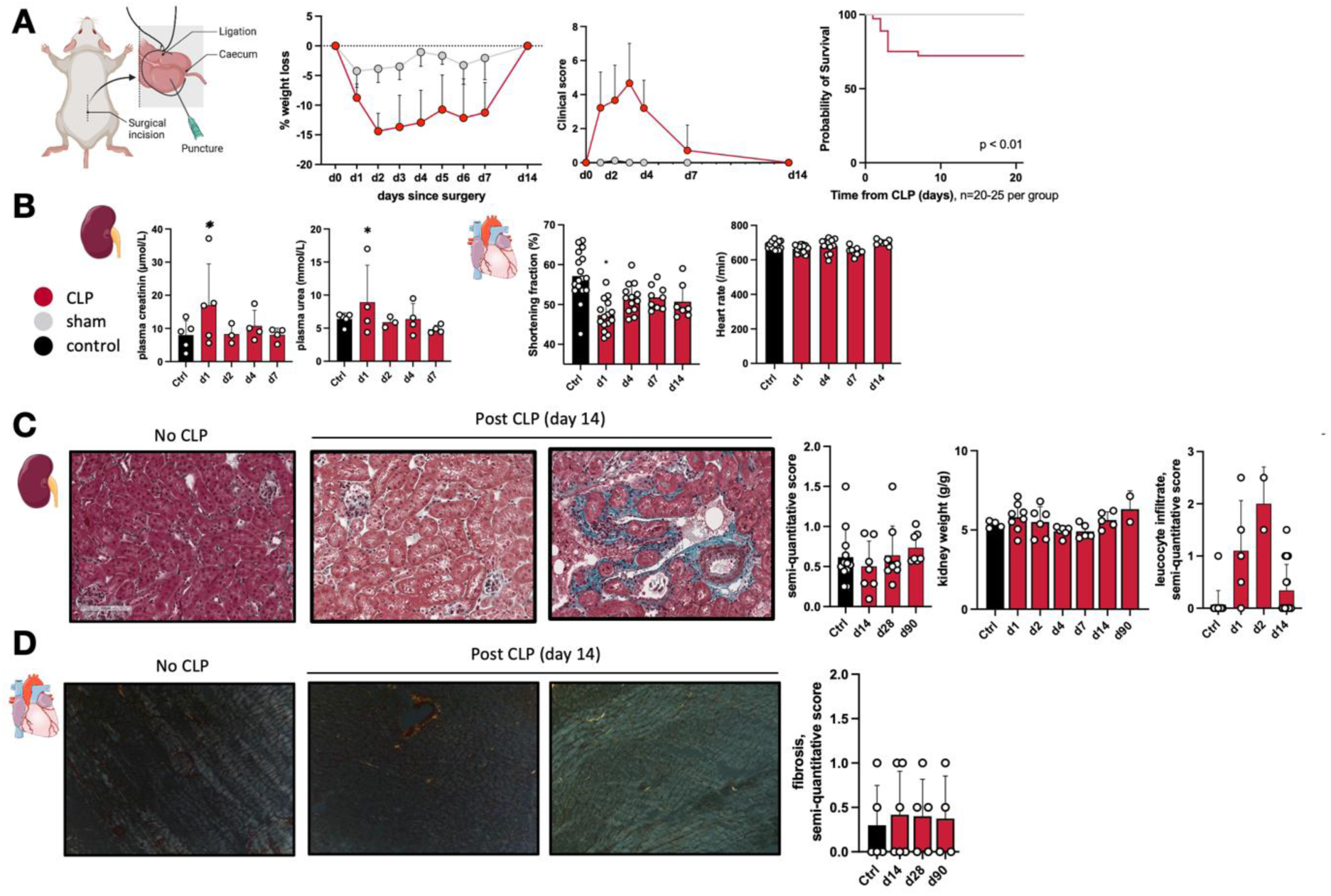
Sepsis is associated with transient and reversible organ dysfunction. **(A)** Diagram of sub-lethal Cecal ligature and punction (CLP) model. Weight loss and behavioral changes were returned to normal values among survivors by day 14. Survival curves confirmed the sub-lethal nature of our model. **(B)** Diagram of plasma urea and creatinine following sepsis as well as echocardiographic constatations in the kidney and heart. **(C)** Histology analyses (Masson Trichrome) of kidney after CLP: fibrosis was assessed using semi-quantitative score by blinded physician, tubular atrophy was assessed by monitoring kidney weights at various timepoints. **(D)** Histology analyses (Sirius red) of cardiac fibrosis after sepsis: fibrosis was assessed using semi-quantitative score by blinded physician. (*p < 0.05; n = 2-10, one-way ANOVA followed by Tukey’s multiple comparisons test). Data are mean ± SEM.

During the acute phase of sepsis, mice developed transient multiorgan failure. Acute kidney injury (AKI) was confirmed by elevated serum urea and creatinine levels, which peaked on day 1 and returned to baseline from day 2 onward (**Figure 1B**). Those changes were transient and not associated with hallmarks of tissue fibrosis: histologic analyses revealed no evidence of renal fibrosis up to 90 days after sepsis, although marked leukocytic infiltration was observed and persisted until day 14. Analysis of kidney weights, used as a surrogate marker of tubular atrophy (14), remained unchanged throughout the 90-day follow-up.

Septic mice also developed sepsis-induced cardiomyopathy (**Figure 1C**), evidenced by a transient decrease in shortening fraction on echocardiography without changes in heart rate on day 1. Cardiac function normalized by day 4. Histologic analysis with Sirius Red staining showed no myocardial fibrosis up to day 90. Taken together, these findings indicate that sepsis causes transient and reversible organ failure but is not, by itself, sufficient to induce long-term fibrosis at the organ level.

### Sepsis induces profound and durable alterations of intra-organ macrophage number and subsets

To investigate how sepsis influences long-term organ function, we examined the dynamics of tissue macrophages after sepsis. Tissue-resident macrophages are key mediators of repair after acute injury in multiple organs (15–18). We therefore quantified macrophage abundance and subsets in the heart and kidney over time following CLP (**Figure 2 and Supplementary Figure 1**).

**Figure 2.**
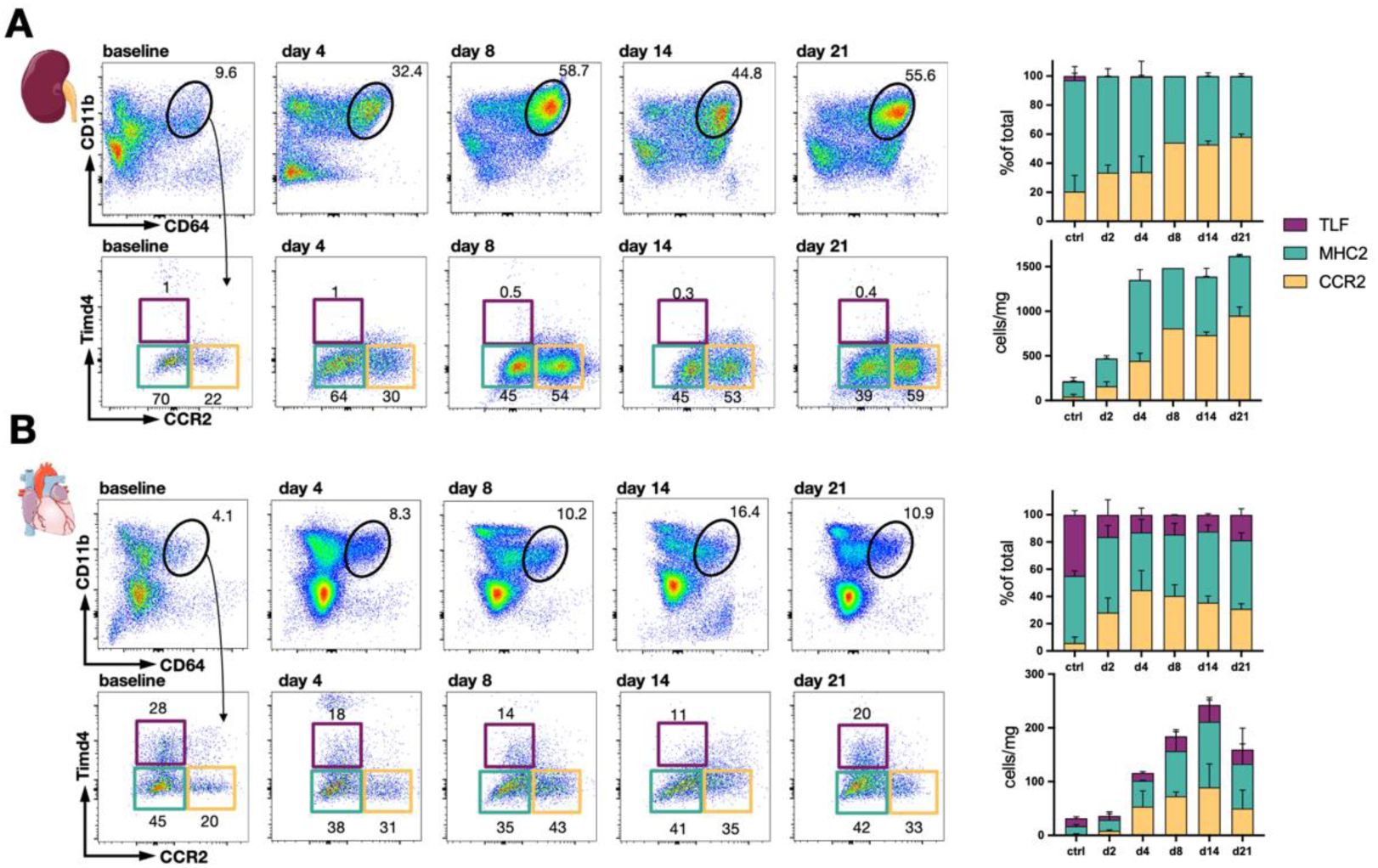
Sepsis induces profound and durable alterations of intra-organ macrophages number and subsets. Kidney **(A)** and heart **(B)** tissues were isolated from adult wild-type mice, and single-cell suspensions were prepared for flow cytometry. Macrophages were identified as live^+^lin^−^CD45^+^CD64^+^CD11b^+^ and then later stratified based on expression of TIMD4 and CCR2. For complete gating strategy see **Supplementary Figure S1.** *p < 0.05; n=2-6; Data are mean ± SEM.

Recent work by Dick et al. identified three conserved subsets of tissue macrophages across species: TLF^+^ (Timd4^+^Lyve1^+^FolR2^+^), CCR2^+^ (Timd4^−^CCR2^+^MHC2^−^) and MHC2^hi^ (CCR2^−^Timd4^−^MHC2^hi^) macrophages (19). These populations can be distinguished by flow cytometry using Timd4 and CCR2 expression as primary gating markers. TLF⁺ macrophages represent a self-renewing, embryonically derived population originating from the yolk sac and fetal liver, while MHC2^hi^ macrophages also arise from embryonic precursors but display distinct transcriptional and surface profiles. In contrast, monocyte-derived macrophages infiltrate tissues with a CCR2⁺ transcriptional program (20).

Using this classification, we characterized the temporal evolution of cardiac and renal macrophage subsets after sepsis.

In the kidney at steady state, tissue macrophages consisted predominantly of the MHC2^hi^ subset, with a smaller proportion of CCR2⁺ macrophages (approximately 20%) and very few TLF⁺ cells (<1%), consistent with previous reports (19). As described by Salei *et al.*, in mice, renal TLF⁺ macrophages are replaced by phenotypically similar cells within a few weeks after birth (21). Following sepsis, we observed a marked increase in total kidney macrophages beginning on day 2 which persisted up to day 21. Phenotypic analysis revealed an almost complete loss of the TLF⁺ subset (**Figure 2A**). The early post-sepsis expansion was driven primarily by CCR2⁺ macrophages, followed by concurrent expansion of both CCR2⁺ and MHC2^hi^ populations. Whereas MHC2^hi^ macrophages predominated at baseline, CCR2⁺ cells became the dominant subset during the subacute and recovery phases (**Figure 2A**).

In the heart, tissue macrophage distribution at baseline differed from that observed in the kidney. The heart retains a substantial pool of yolk sac– and fetal liver–derived macrophages with self-renewing capacity(22). At steady state, TLF⁺ macrophages accounted for 30–40% of the total macrophage population, while MHC2^hi^ macrophages represented approximately 50%, and CCR2⁺ macrophages only 5–10% (**Figure 2B**). Following sepsis, the total number of cardiac macrophages markedly increased, though this expansion occurred later than in the kidney, beginning around day 4, and declined after day 14. The CCR2⁺ subset showed a transient rise, before returning toward baseline. The proportion of TLF⁺ macrophages decreased during the early phase but stabilized thereafter, with a delayed increase in absolute numbers after day 4. In contrast, the relative abundance of MHC2^hi^ macrophages remained stable over time. These findings suggest either differentiation of monocyte-derived macrophages or expansion of the embryonically derived TLF⁺ pool. While Dick *et al.* previously demonstrated that this subset can self-renew in adulthood with minimal hematopoietic contribution, such dynamics have not been reported in a septic context (22).

Taken together, these findings indicate that, despite apparent recovery from acute organ failure, sepsis induces profound and long-lasting alterations in tissue macrophage composition. In both the kidney and heart, we observed a sustained increase in CCR2⁺ macrophages. Although previous studies have suggested that these cells are monocyte-derived, this has not been directly demonstrated in the context of sepsis.

### Sepsis induces intense and sustained hematopoietic response arising from both bone marrow and splenic reservoirs

Emergency hematopoiesis is a key feature of the early inflammatory response, and previous work has identified two distinct waves of monocyte release following sepsis onset (23). To assess hematopoietic dynamics over time, we analyzed bone marrow and splenic responses after CLP. The spleen is known to play a central role in emergency hematopoiesis during sepsis (24). We quantified lineage⁻ cKit⁺ Sca-1⁺ (LSK) progenitor cells in both bone marrow and spleen (**Figure 3 and Supplementary Figure 1**). Sepsis induced a modest expansion of LSK progenitors in the bone marrow but a pronounced increase in the spleen, peaking at day 14. Similarly, downstream granulocyte–macrophage progenitors (GMPs) and myeloid-derived progenitors (MDPs) increased markedly from day 2 onward, with sustained myelopoietic activity in spleen compartments up to day 14 while it tended to decrease afterwards.

**Figure 3.**
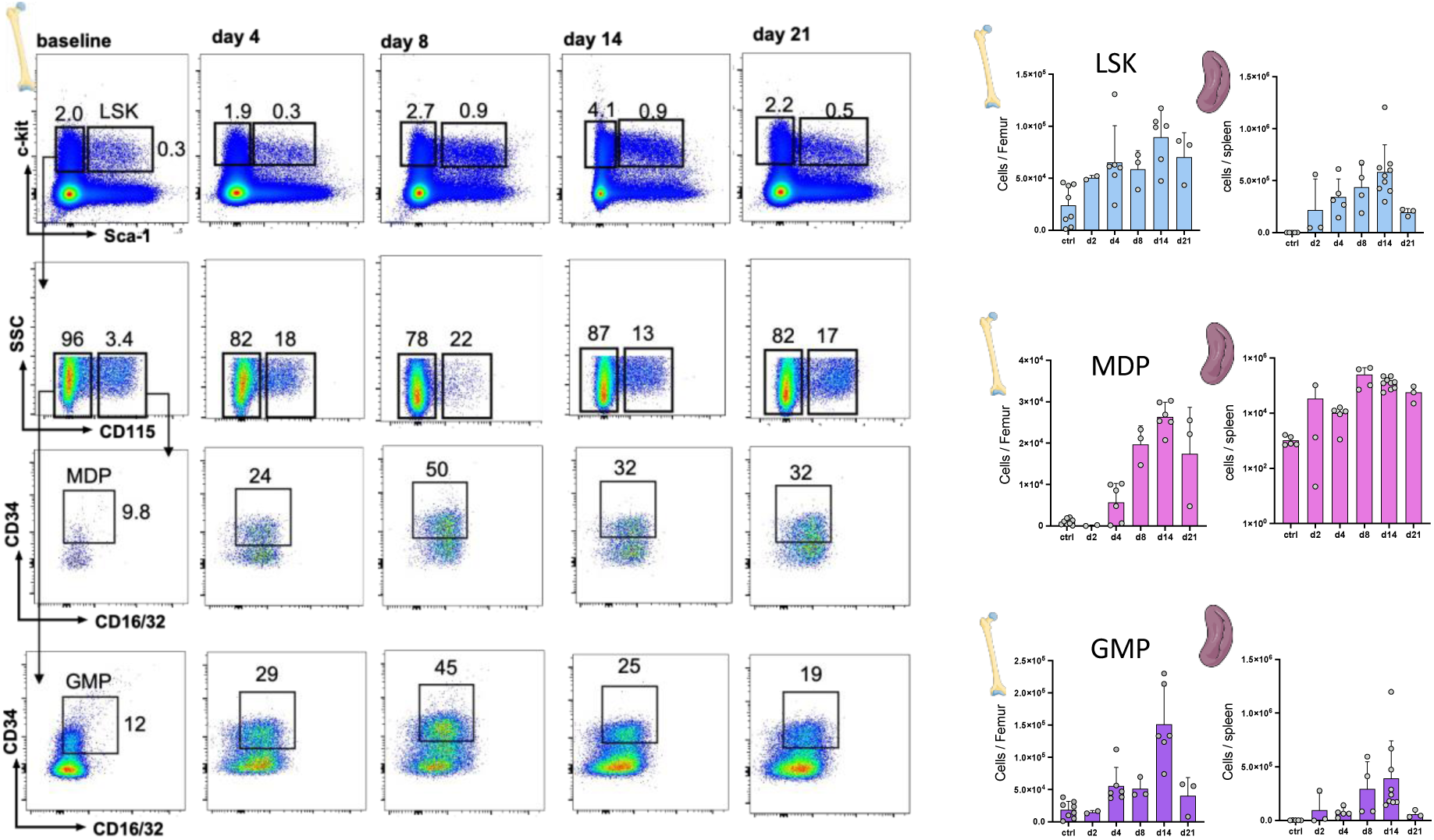
Sepsis induces intense and sustained hematopoietic response arising from both bone marrow and splenic reservoirs. Flow cytometric quantification of hematopoietic precursors in bone marrow and spleen following sepsis. LSK were gated based on expression of live^+^lin^−^sca1^+^ckit^+^. GMP were gated on live+lin^−^ckit+sca1^−^CD115^−^CD16/32^+^CD34^+^ and MDP were gated based on live^+^lin^−^ckit^+^sca1^−^CD115^+^CD16/32^+^CD34^+^. For complete gating strategy of the spleen, see **Supplementary Figure S1.** *p < 0.05; n=2-6; Data are mean ± SEM.

These findings indicate that, despite apparent resolution of clinical symptoms, bone marrow and splenic reservoirs contribute to a sustained hematopoiesis after sepsis. This prolonged hematopoietic activation persists for weeks and may be responsible for the observed increment in tissue macrophages we observed.

### Fate-Mapping Analysis Identifies Monocyte-Derived Macrophages as the Main Source of Tissue Macrophage Expansion after Sepsis

The marked expansion of tissue macrophages after sepsis raises the question of their origin, whether these cells arise from self-expansion of resident macrophages or from the recruitment of circulating monocytes that differentiate into macrophages. Because both macrophage number and phenotype evolved over time, we sought to delineate the relative contribution of each population.

To this end, we used using tamoxifen inducible Cx3cr1CreER/^+^R26tdTomato/^+^ fate-mapping model (7) as described by Kretzschmar et al. (25) (see **method section** and **Supplementary figure 2** for detail). This model takes advantage of the long lifespan of macrophages, which express CX3CR1, compared to the short life of circulating monocytes, whose progenitors do not express CX3CR1. Prior studies have shown robust induction of reporter gene expression in CX3CR1-expressing resident macrophages, and transient labeling of blood monocytes (12,19,26). Six- to ten-week-old c57Bl6 mice were fed using tamoxifen-containing chow for 10 days (**Figure 4A**). By day 10, complete Cre recombinase induction was confirmed by flow cytometry, with 100% Tomato (Tdt) retention in blood monocytes and tissue macrophages of the kidney and heart. (**Figure 4B**.).

**Figure 4.**
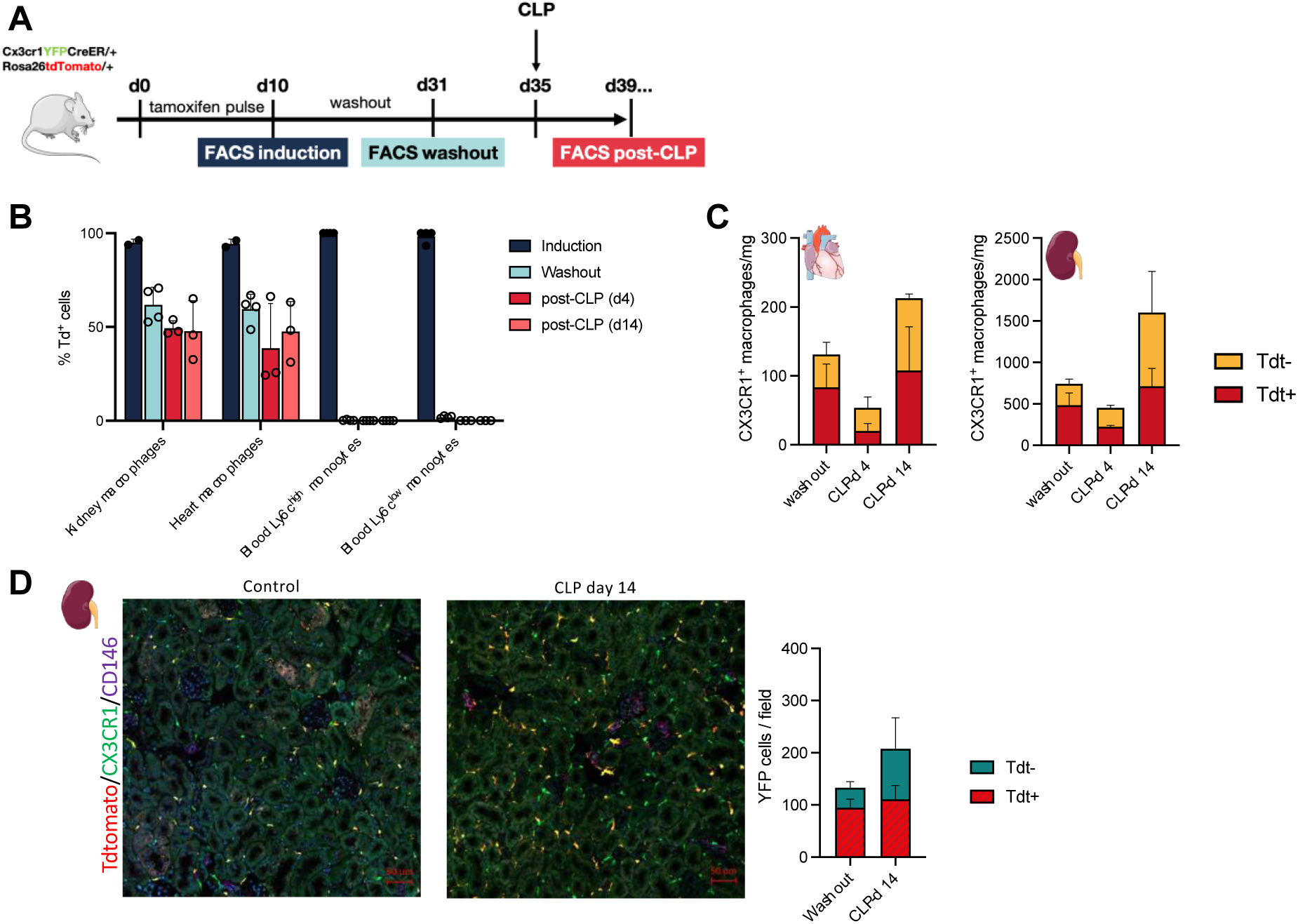
Fate mapping analysis identifies monocytes-derived macrophages as the main source of tissue macrophages expansion during sepsis. **(A)** 6–10-week-old c57Bl6 Cx3Cr1YFPCreER/+Rosa26Tdtomato/+ mice were fed with tamoxifen-containing chow for 10 days. By day 10, we confirmed by flow-cytometry the complete induction of Cre Recombinase. After a period of washout of 21days (confirmed by flow cytometry), mice were subjected to cecal ligature punction. Flow-cytometry was used at different timepoints to assess abundance and Tdt+ retention of intra-organ macrophages. **(B)** Percentage of Td+ cells in tLy6C^hi^ monocytes, Ly6C^lo^ monocytes, kidney and heart after induction, after washout or at different timepoints following sepsis. **(C)** For each timepoints, total number of intra-organ macrophages was assessed and Tdt+ retention was computed. **(D)** Representative immunofluorescence staining of EYFP (green), Tdtomato (red) and CD146 (violet) in of adult Cx3Cr1YFPCreER/+Rosa26Tdtomato/+ mice at day 14 after sham or CLP, x20 magnification, Quantification of the proportion of Tdt+ in macrophages was performed by blinded physician (n=2-3).

After a 21-day washout period, Tdt expression disappeared entirely from circulating monocytes, indicating full renewal of the blood monocyte pool, while persisting in tissue macrophages (**Figure 4B**). Thus, in this model, **Tdt⁺** cells represent long-lived, tissue-resident macrophages, whereas **Tdt⁻** cells derive from newly recruited monocytes. Notably, even under baseline conditions, 30–40% of kidney and cardiac macrophages were replaced by circulating monocytes within 21 days, consistent with previous observations by Dick *et al* (19).

Following sepsis, we observed a transient decrease in Tdt⁺ macrophages in both kidney and heart at day 4 (**Figure 4C**), indicating an early loss of resident macrophages consistent with the “disappearance reaction” previously reported in models of viral or *Listeria* infection (27,28(29)) but not yet described in sepsis. This loss was followed by a progressive expansion of the Tdt⁺ population by day 14, reflecting self-renewal of surviving resident macrophages, as Tdt labeling is stably transmitted to daughter cells (30).These findings align with prior work demonstrating the proliferative potential of intra-organ macrophages (31,32), here extended to the septic context.

In parallel, we observed a pronounced recruitment of Tdt⁻ monocyte-derived macrophages between days 4 and 14, accounting for most of the overall increase in macrophage numbers as the resident pool gradually returned to baseline (**Figure 4C**). Immunofluorescence imaging of kidney sections from fate-mapped mice confirmed these findings (**Figure 4D**).

Collectively, these results demonstrate that macrophage dynamics after sepsis involve both self-renewal of tissue-resident macrophages and sustained recruitment of monocyte-derived macrophages. The latter predominates during the subacute phase and parallels the activation of emergency hematopoiesis, whereas resident macrophage proliferation occurs later in the recovery period.

### Similar immune populations colonize the heart and kidneys after sepsis

The persistence of tissue macrophages after sepsis suggested potential remodeling of their transcriptional landscape, affecting both resident and monocyte-derived subsets. To define these changes, we analyzed the transcriptomic profiles of intra-organ myeloid cells from sham controls and from mice 14 days after sepsis, a time point when clinical recovery was complete. Myeloid cells (live⁺CD45⁺CD11b⁺) were sorted from kidney and heart (n = 3 per condition, pooled before sequencing) (**Figure 5A**). After exclusion of low-quality cells, lymphocytes, and neutrophils (**Supplementary Figures 3**), 7,642 kidney cells and 963 heart cells passed quality control for downstream analysis.

**Figure 5.**
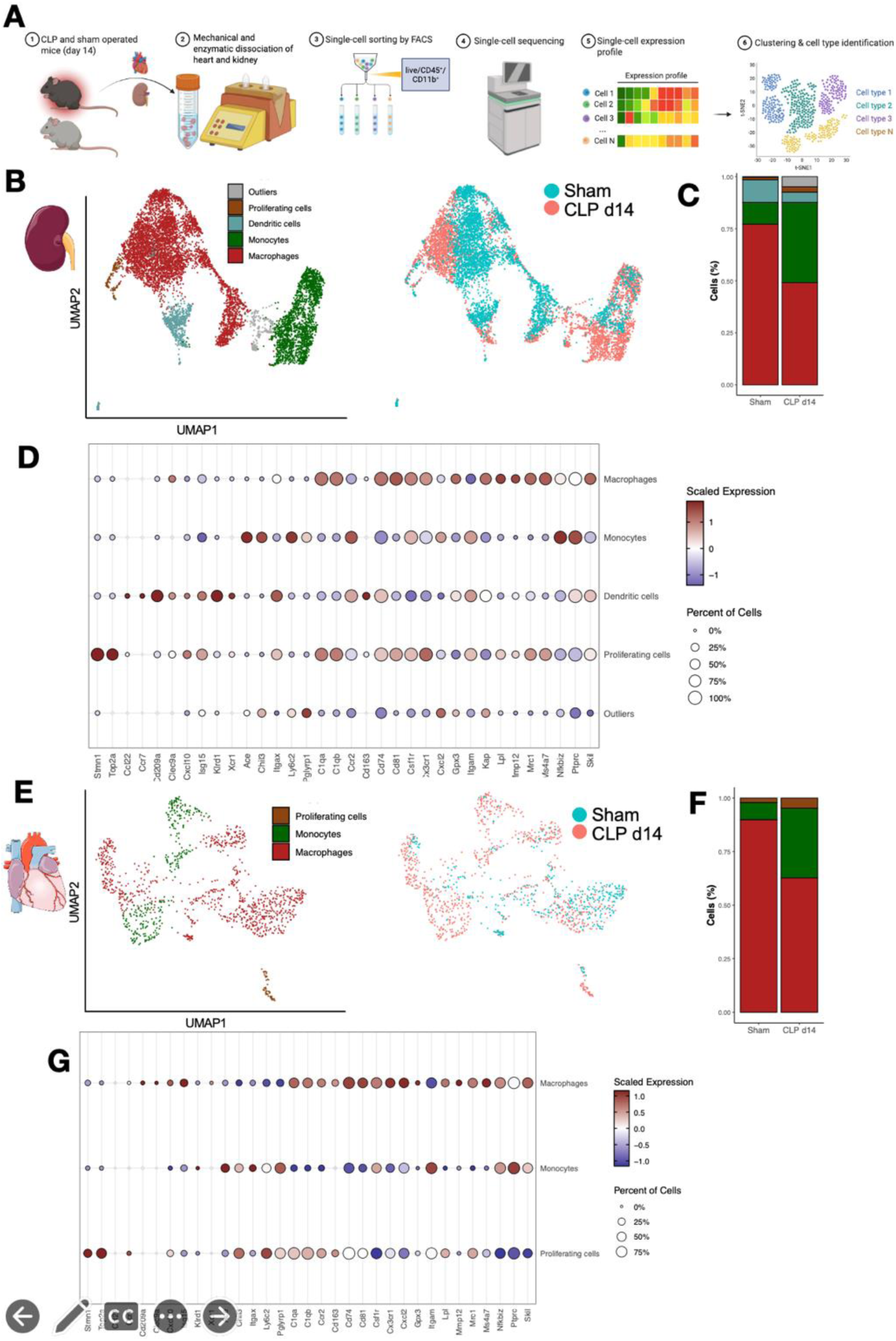
Similar immune populations colonize the heart and kidney after sepsis. **(A)** Experimental design: myeloid cells (live⁺CD45⁺CD11b⁺) were sorted from kidney and heart (n = 3 per condition, pooled before sequencing) from mice on day 14 after CLP and sham operated mice. **(B, E)** UMAP representation of unsupervised clustering of intra-organ myeloid cells (live⁺CD45⁺CD11b⁺) from kidney (B) and heart (E) at baseline (sham) and 14 days after CLP-induced sepsis. (**C, F)** Relative proportion of identified myeloid populations in kidney (C) and heart (F) comparing sham and septic mice. **(D, G)** Expression heatmaps of canonical marker genes used for cell-type annotation across clusters. Data are pooled from n = 3 mice per condition. **Abbreviations:** CLP = cecal ligation and puncture; UMAP = uniform manifold approximation and projection.

Unsupervised clustering revealed five distinct myeloid populations in the kidney and three in the heart (**Figures 5B and 5E**). Cell identities were assigned using canonical marker genes for macrophages (*C1qa, C1qb, Ccr2, Cd163, Cd74, Cd81, Csf1r, Cx3cr1, Cxcl2, Gpx3, Itgam, Lpl, Mmp12, Mrc1, Ms4a7, Nfkbiz, Ptprc,* and *Skil*), monocytes (*Ace, Chil3, Itgax, Ly6c2, Pglyrp1*), dendritic cells (*Ccl2, Ccr7, Cd209a, Clec9a, Cxcl10, Isg15, Klrd1, Xcr1*), and proliferating cells (*Stmn1, Top2a*) (**Figures 5D and 5G, Supplementary Table 1**).

Compared with sham controls, kidneys and hearts harvested on day 14 after CLP displayed a shift in immune cell composition, characterized by an increased proportion of monocytes in both organs (**Figures 5C and 5F**). These results are consistent with our earlier analyses, which showed a higher number of macrophage-rich regions in post-septic tissues relative to controls.

### Sepsis Triggers Accumulation and Differentiation of Profibrotic Monocytes in the Kidney and Heart

We next focused our analysis on monocyte populations. Two transcriptionally distinct monocyte subsets were identified in both kidney and heart (**Figures 6A and 6D**). Sub-clusters were annotated based on established markers of monocyte heterogeneity in health and disease (20,33–36). One subset exhibited high expression of *Cx3cr1*, *Ace*, and *Pglyrp1*, consistent with patrolling monocytes (**Figure 6B**) (*33*). The second subset expressed *Ly6c2* and *Chil3*, identifying them as Ly6C2⁺ monocytes. As patrolling monocytes arise exclusively from Ly6C⁺ precursors (7,37), both subsets likely originate from initially recruited Ly6C⁺ monocytes.

**Figure 6.**
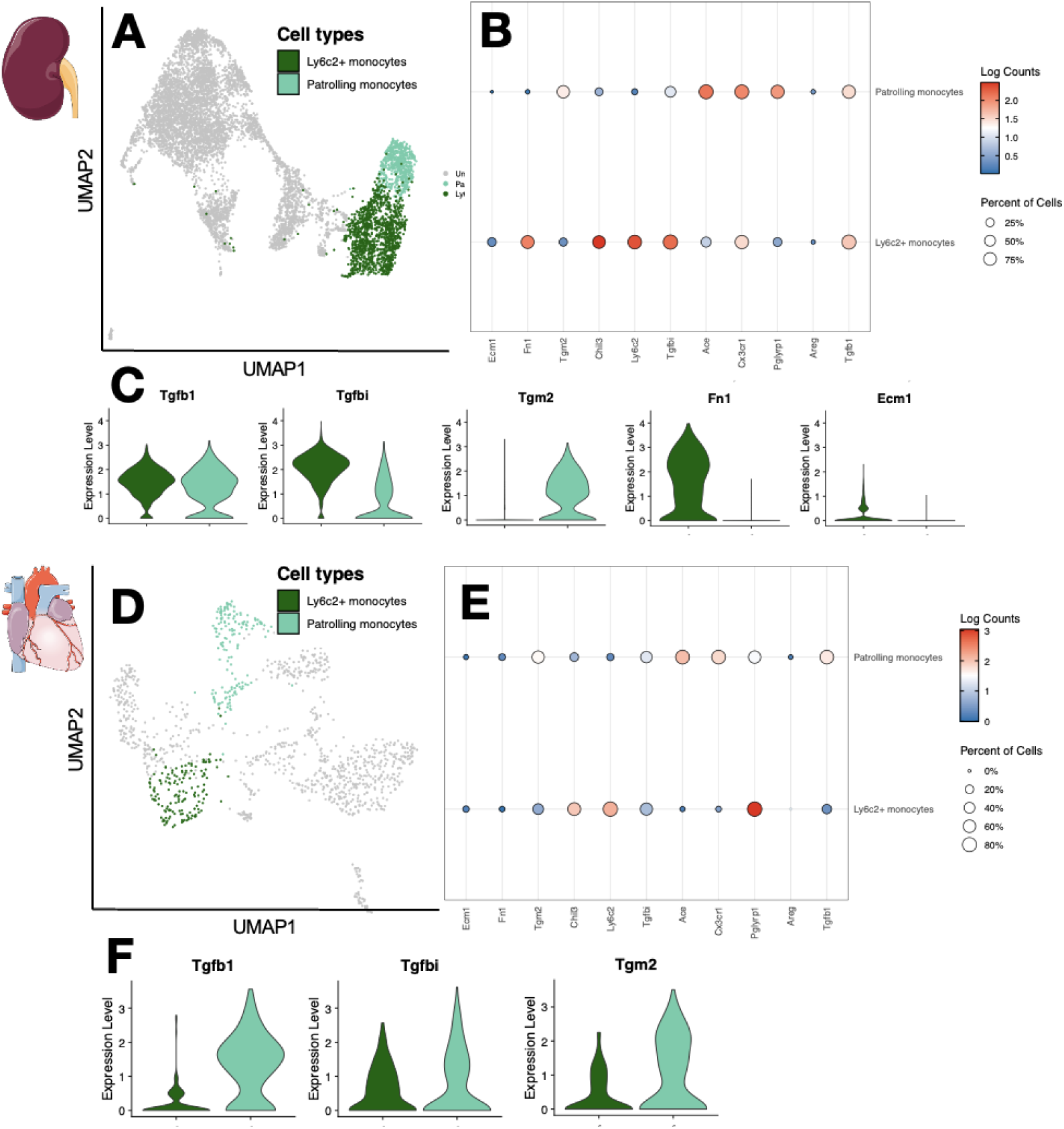
Sepsis triggers accumulation of profibrotic monocytes in the kidney and heart. **(A, F)** UMAP visualization of monocyte clusters from kidney (A) and heart (D) at day 14 after CLP, identifying Ly6C2⁺ and patrolling (Cx3cr1⁺) monocyte subsets. **(B, E)** Feature plots showing expression of key subset-defining genes. **(C, F)** Violin plots highlighting organ-specific enrichment of profirbrotic (*Tgbi*, *Tgfb1*), extracellular-matrix (ECM) genes (*Fn1*, *Ecm1*) and cross-linkers (*Tgm2*). **Abbreviations:** CLP = cecal ligation and puncture; ECM = extracellular matrix; GSEA = gene-set enrichment analysis; UMAP = uniform manifold approximation and projection.

While the proportion of two monocytes population tended to maintain, absolute number of monocytes greatly increased after CLP (**Figure 5B**). Ly6c2+ monocytes harbored high expression of pro-fibrotic genes such as *Tgfb*1 and *Tgfbi* in the kidney (**Figures 6C**), while these genes were more expressed in the patrolling monocytes subset (**Figure 6F)**. *Ly6C2*^+^ monocytes expressed gene of extracellular matrix (ECM) component *(Ecm1)* in the kidneys (**Figure 6C 6F**), while patrolling monocytes from both organs harbored high expression of ECM crosslinkers (*Tgm2*, **Figures 6C, F**).

### Macrophage sub-populations show heterogeneity after sepsis

We identified 5 macrophages subtypes in the kidney, and 4 in the heart with different phenotypic signatures (**Figures 7A, D**). Identification of sub-populations was made on the basis of previously published literature on tissue macrophages heterogeneity in health and disease (20,33–36) (**Supplementary Table 1**).

**Figure 7.**
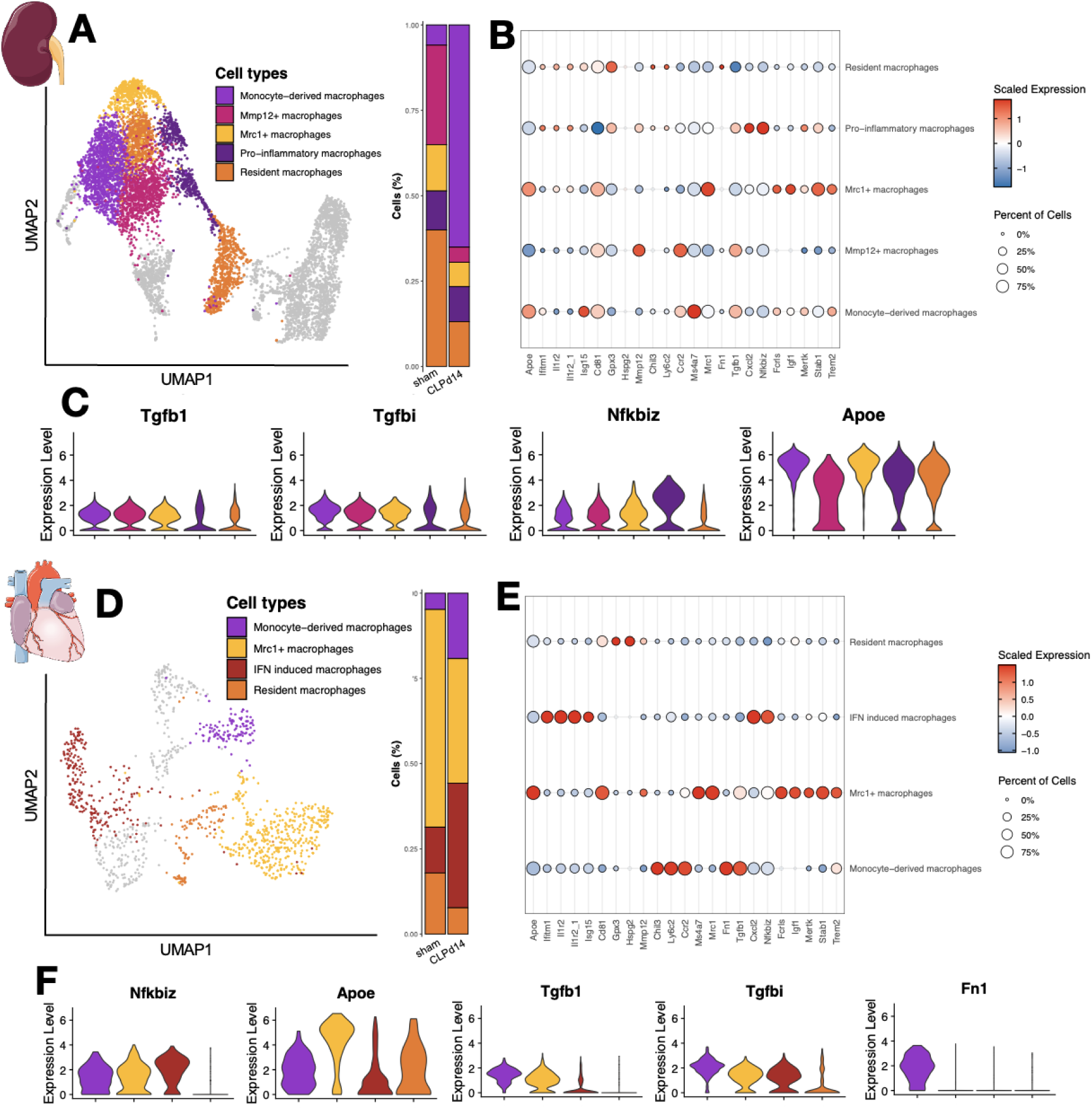
Sepsis remodels the macrophage landscape in kidney and heart. **(A, D)** UMAP plots of macrophage subsets identified in kidney (A) and heart (D) at day 14 after CLP compared with sham controls, showing five and four clusters respectively. Quantification of macrophage subset proportions in sham and septic mice showing reduction of resident and Mrc1⁺ macrophages and increase of CCR2⁺ and IFN-induced populations after sepsis. **(B, E)** Heatmaps of representative marker genes defining macrophage subtypes: resident (*Gpx3, Hspg2*), proinflammatory (*Nfkbiz, Cxcl2*), reparative (*Mrc1, Apoe, Igf1, Trem2, Mertk*), monocyte-derived (*Ccr2, Ms4a7, Chil3, Ly6c2, Fn1, Tgfb1*), and Mmp12⁺ or IFN-induced (*Il1r2, Isg15*) macrophages. (**C, F**) Violin plots highlighting organ-specific enrichment in inflammatory (*Nfkbiz*), pro-fibrotic (*Tgbi, Tgb1, Fn1*) and decrease in regenerative (*Apoe*) genes. Data are pooled from n = 3 mice per condition. **Abbreviations:** CLP = cecal ligation and puncture; UMAP = uniform manifold approximation and projection.

In the kidney, one subset corresponded to **resident macrophages**, characterized by high expression of **Gpx3**, encoding glutathione peroxidase 3, an enzyme that promotes hydrogen peroxide metabolism and protects cell membranes from oxidative injury (42). A comparable resident subset was identified in the heart, but these cells also expressed **Hspg2** (*Perlecan*), a proteoglycan involved in extracellular matrix stabilization (41) (**Figure 7B, E**).

A second macrophage population expressed high levels of **Nfkbiz** and **Cxcl2**, a chemokine critical for monocyte recruitment in the injured kidney (43). This subset likely contributes to sustained monocyte infiltration after sepsis and was designated as **proinflammatory macrophages**. In the heat, those cells also expressed high level cytokine receptor *Il1r2* and interferon-stimulated genes (*Isg15*, *Ifitm1*) and were therefore labelled **Interferon-induced macrophages (Figure 7F)**.

A third subset, defined by strong expression of **Mrc1** (encoding the mannose receptor), exhibited a reparative transcriptional profile with enrichment in scavenger receptors (*Mrc1*, *Fcrls*, *Stab1*) and genes involved in anti-inflammatory and regenerative functions, including *Apoe*, *Igf1*, *Trem2*, and *Mertk*. These features are consistent with a reparative macrophage phenotype. Notably, TREM2^hi^ macrophages have been shown to protect the septic heart by clearing damaged cardiomyocyte mitochondria; loss of Trem2 exacerbates cardiac dysfunction (34).

Lastly, we identified a **Mmp12⁺** macrophage subset in the kidney, abundant in control mice but nearly absent after sepsis (**Figure 7B**). **Mmp12**, a macrophage-specific metalloproteinase, is implicated in matrix remodeling and repair, and its expression marks a reparative phenotype in models of liver fibrosis.

Quantitative analysis revealed a decrease in the proportions of resident and Mrc1⁺ macrophages in septic mice compared with controls in both kidney and heart (**Figures 7B, E**). In the kidney, Mmp12⁺ macrophages also declined, whereas CCR2⁺ monocyte-derived macrophages markedly increased. Similarly, in the heart, sepsis induced an expansion of IFN-induced and CCR2⁺ macrophages (**Figure 7D**).

Taken together, these findings demonstrate that sepsis profoundly remodels the macrophage landscape in both kidney and heart, characterized by a loss of resident and reparative macrophages and the expansion of proinflammatory and monocyte-derived subsets.

### Sepsis Impairs Organ Repair after a Secondary Insult

Building on our previous findings, sepsis induces transient organ failure and long-term remodeling of the macrophage network, characterized by the recruitment of monocyte-derived, proinflammatory, and profibrotic populations. The functional consequences of these immune alterations on tissue repair remained to be determined.

To address this question, we developed a clinically relevant double-hit model. Mice underwent either sham surgery or sublethal CLP, followed 14 days later by implantation of a micro-osmotic pump delivering angiotensin II and exposure to 2% NaCl drinking water (**Figure 8A**). This approach recapitulates a common post-septic phenotype characterized by persistent activation of the renin–angiotensin system and accelerated cardio-renal disease.

**Figure 8.**
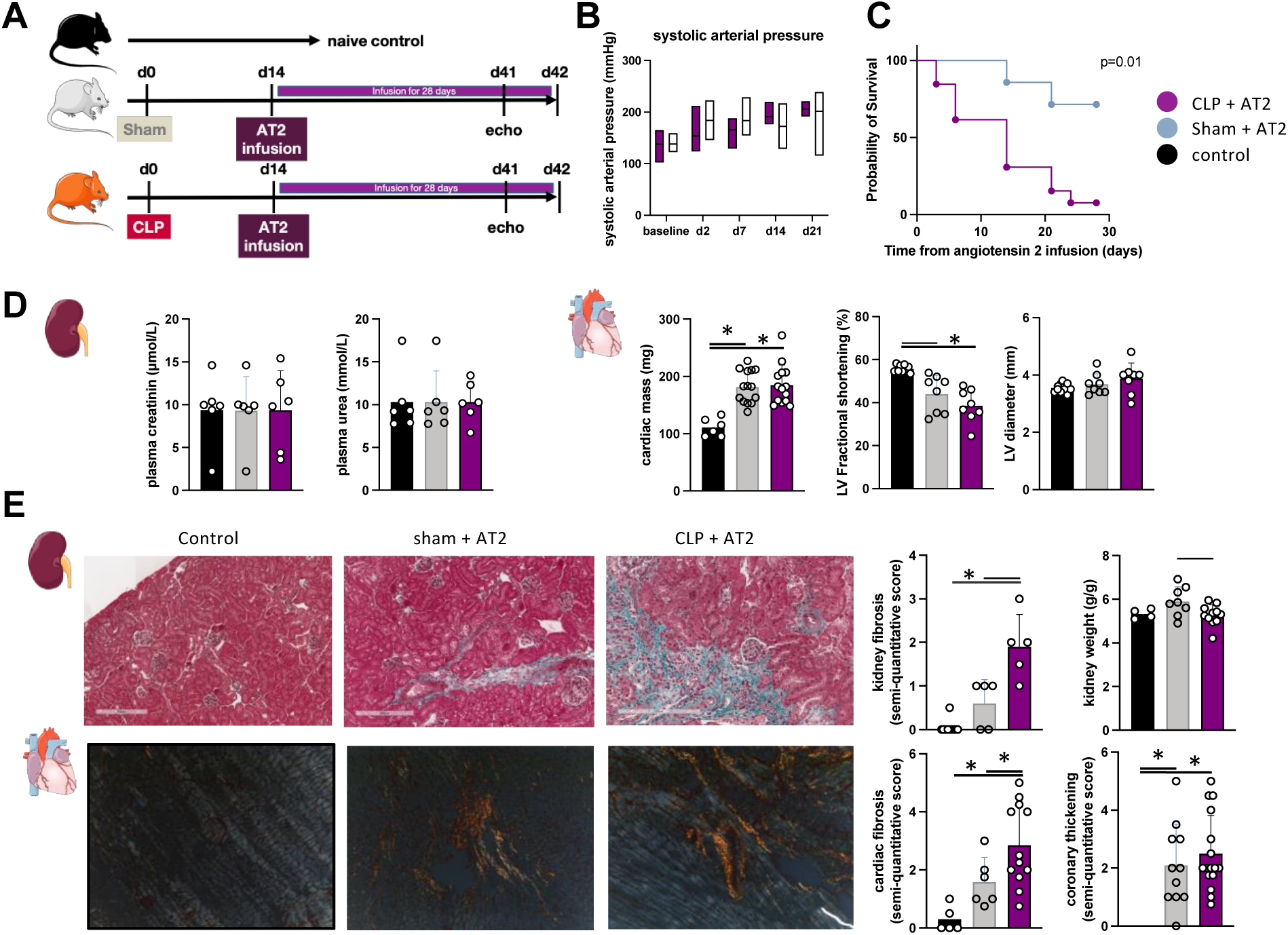
Sepsis causes altered organ repair after second hit. **(A)** C57bl6 WT mice were randomly assigned to receiver either CLP or sham operation. After complete recovery from the septic episode (day 14), sepsis-surviving mice and sham operated mice were subjected to continuous angiotensin (AT) 2 infusion (800ng/kg/min) combined with salty drinking water (NaCl 0.2%) for 28 days. Echocardiographic evaluation was performed by blinded physician just before sacrifice. **(B)** AT 2 infusion induced hypertension as confirmed by arterial pressure monitoring, appearing from day 2 **(C)** Survival curves of WT mice when exposed to AT2 demonstrates an important mortality among sepsis survivor mice when exposed to AT 2 as compared to sham operated mice **(D)** Plasma urea and creatinine did not differ with baseline after AT 2 infusion. Echocardiography revealed increased cardiac mass and reduction of fractional shortening consistent with AT 2 induced cardiac hypertrophy **(E)** Weighting of kidney was in favor of tubular atrophy in CLP mice after AT2 infusion. Histological analyses of kidney (Masson trichrome) revealed fibrosis in both conditions, with a strong increase among sepsis-surviving mice. Heart analyzes (Sirius red) revealed coronary thickening, consistent with intense sheer stress induced by AT 2 infusion. Fibrosis was present in both conditions; it was more important in sepsis survivors as compared with sham operated mice. Coronary thickening and fibrosis were assessed using semi-quantitative score by a blinded physician; t test was performed; *p<0.05; **p<0.01; data are mean+/− SEM.

Angiotensin II infusion induced comparable hypertension in both sham and CLP-operated mice (**Figure 8B**). However, 28-day mortality was markedly higher in previously septic animals (p < 0.01; **Figure 8C**). No aneurysm rupture or intracranial hemorrhage was detected at necropsy.

At the organ level, classical kidney function biomarkers remained unchanged between groups (**Figure 8D**), but kidney weight was significantly reduced in CLP mice (**Figure 8E**). Angiotensin II infusion led to increased cardiac mass and reduced contractility in both groups, with no significant difference in systolic parameters (**Figure 8D**). Notably, histologic analyses revealed markedly enhanced cardiac and renal fibrosis in post-septic mice compared with controls at day 28 after angiotensin II exposure (**Figure 8F**).

These findings demonstrate that while sepsis alone does not induce long-term fibrosis, it profoundly alters organ resilience, predisposing tissues to maladaptive repair after subsequent injury. Given the central role of macrophages in tissue regeneration, this defective repair phenotype may reflects the post-septic reorganization of macrophage populations toward a profibrotic and inflammatory profile. Together, these results provide a mechanistic explanation for the increased risk of chronic organ dysfunction observed in sepsis survivors, consistent with clinical trajectories of recurrent tissue injury and progressive cardio-renal decline.

## DISCUSSION

Awareness of the long-term consequences of sepsis has increased worldwide. Sepsis remains a frequent and severe condition with high short-term mortality; however, mounting evidence suggests that much of the morbidity and mortality attributable to sepsis occurs in the months and years following ICU discharge (48). Sepsis survivors experience a high rate of rehospitalization and are at increased risk of developing chronic kidney disease and heart failure (4, 5, 49–51). Despite this, the mechanisms linking an acute septic event to chronic organ dysfunction remain poorly understood.

In this study, we demonstrate that sepsis induces transient, reversible organ failure but also causes long-lasting reorganization of tissue macrophage populations. We show that sepsis alters the transcriptional landscape of tissue-resident macrophages and promotes the recruitment of monocyte-derived macrophages with proinflammatory and profibrotic profiles. These changes create a local immune environment unfavorable to normal tissue repair, rendering organs more susceptible to maladaptive repair after a secondary insult. Collectively, our findings suggest that sepsis leaves behind a durable “immune scar” that compromises organ resilience and may contribute to the development of chronic organ failure.

Sepsis, cardiovascular diseases and chronic kidney failure have been linked by numerous observational studies (4,5). The occurrence of acute kidney injury during sepsis hospitalization was a major risk factor to develop CKD. Heart and kidney both recruit thousands of circulating myeloid cells during sepsis, those cells seem to have a protective role in the acute phase, protecting the septic organs from remote injury (52). However, switching phenotype is crucial for monocyte-derived macrophage to favor healing versus fibrosis in various contexts (16,53,54) failure to do so may be responsible for the maladaptive repair observed in various condition (55). In this perspective, we hypothesized that a second hit occurring before complete intra-organ immune restoration (*immune reset*), but after apparent recovery, might favor emergence of fibrosis and tissue atrophy over tissue regeneration.

To explore this, we established a clinically relevant *two-hit* model combining sublethal sepsis (CLP) followed by angiotensin II infusion—a condition that mimics the hyper-angiotensin state frequently observed in sepsis survivors and associated with accelerated cardio-renal decline (56–58). Hypertension is very frequent among sepsis patients and is associated with worse long-term outcome (59). Angiotensin 2 infusion is a well described model which causes robust hypertension and which is pro-fibrotic by itself (60,61). This new two-hit model can be used as a readout to investigate long-term sepsis consequences on organ function and repair by generating, as seen in clinical practice, multiple hits after initial sepsis insult. Interestingly, macrophage in the kidney had high expression of angiotensin receptor associated protein (A*gtrap)*, which suggest they might be sensible to angiotensin-2 infusion.

Sepsis by itself did not induce heart nor kidney fibrosis in our model up to 3 months, but was responsible for long term intra-organ modification of the immune landscape with profound modification of number and phenotype of intra-organ macrophages. We here demonstrate that monocyte-derived-macrophages are responsible for the global increment in intra-organ macrophage pool, which lasts for several weeks despite complete recovery. We also confirmed that monocyte-derived macrophage bears a pro-fibrotic, pro-inflammatory phenotype as compared to resident macrophages.

In summary, sepsis has a profound impact on tissue macrophages populations especially through the recruitment of pro-fibrotic monocyte-derived macrophages that are associated with altered organ repair. Macrophages populations and function could therefore represent a new therapeutic target to treat the long-term consequences of sepsis.

## MATERIAL AND METHODS

### Mice

C57BL/6J mice were purchased from Janvier Laboratory. RosaTd reporter mice (Jackson Laboratory, #007914) were crossed with Cx3cr1CreERT2 (Jackson Laboratory, #020940) for the FateMapping experiments. Recombination in heterozygous Cx3cr1CreER/+R26tdTomato/+. All mice were housed under specific pathogen-free conditions in our animal facility at the Centre Viggo Petersen in Lariboisière University Hospital. For the fate mapping experiments, male and female mice were used due to the rarity of the specimen. Mice were 6-10 weeks old at the start of each experiment. Mice were randomly assigned to the experimental groups. All experiments were approved by our local ethic committee and the Ministère de l’Enseignement et de la Recherche (APAFIS #28236).

### Cecal ligation and puncture surgery

Cecal ligation and puncture was performed in mice anesthetized via intraperitoneal (i.p) injection of Ketamine (100mg/kg) and Xylazine (10mg/kg). An abdominal midline incision was made (0.7cm) and the *linea alba* of the abdominal musculature was dissected. For sub-lethal peritonitis, the cecal tip was exteriorized, and the ligation was performed approximately 10% from its extremity with a 4.0 silk suture. The cecum was punctured with a 24-gauge needle, and a small amount of feces was extruded through the perforation. The cecum was next returned into the peritoneal cavity, and the abdominal musculature was closed with a 4.0 monofilament suture. Mice were treated i.p. with 1mL sterile saline. The skin incision was closed with staples. Sham-operated mice underwent the same procedure except for the caecal ligation and puncture.

### Double hit model: Angiotensin 2 infusion

After a first hit with CLP or Sham surgery, surviving mice were subjected to a second challenge with angioII infusion. AngioII (Sigma-Aldrich) was infused at 800ng/kg/min via a subcutaneous osmotic pump (ALZET®) for 28 days, together with addition of salt in the daily water (NaCl 2%). Diet and tap water were provided *ad libitum* throughout the experiment.

### Echocardiography in mice

Cardiac parameters were measured by transthoracic echocardiography using ACUSON S3000 ™ (Siemens) Cardiovascular Ultrasound system equipped with a 14-MHz linear transducer. Mice were anesthetized by ketamine (80 mg/kg) during measurements. All echocardiography were performed blindly.

### Histochemistry

Kidneys were collected and immersed in 5% formo-acetic acid during 24 h, dehydrated, and embedded in paraffin according to the standard protocols. Sections at 4 μm thickness (HM 355S microtome, Thermo scientific) were stained with masson trichrome coloration. Harvest hearts were fixed with OCT and cryostat sections (7μm) of the ventricles were performed. A sirius red staining of the collagens was realized. All images were acquired on a Leica microscope. Quantification of fibrosis was performed by expert physicians who were blinded of the experiments.

### Tissue dissociation and flow cytometry

Heart and kidney were collected in PBS and processed immediately. Organs were finely chopped, and digested with MACS® tissue dissociation kit 2 (Miltenyi Biotec) using GentleMACSTM Octo Dissociator (Miltenyi Biotec) according to the manufacturer instruction. The spleen was harvested and gently mashed, while the bone marrow was collected via centrifugation of the left femur. The cell suspension from all organs was filtered through a 70µm strainer and red blood cells (RBCs) were lysed in ammonium chloride potassium (ACK) lysis buffer twice for 5min at RT.

For each sample 2 million cells were stained. In heart and kidney, macrophages were identified as live^+^ lin^−^ CD45^+^ CD64^+^ CD11b^+^, and macrophages subpopulation were later stratified based on expression of TIMD4 and CCR2. In bone marrow and spleen, hematopoietic precursors were identified as LSK (live^+^ lin^−^ sca1^+^ ckit^+^), GMP (live+ lin^−^ ckit+ sca1^−^ CD115^−^ CD16/32^+^ CD34^+^) and (live^+^ lin^−^ ckit^+^ sca1^−^ CD115^+^ CD16/32^+^ CD34+). A full description of the antibody used is provided in Supplementary Materiel. Flow cytometry was carried out using a BD® LSR II Flow Cytometer (Becton Dickinson). Blind analysis of the data was performed using FlowJo™ Software (Tree Star).

### Fate mapping

*Heterozygous Cx3cr1*^CreER/+^*R26*^tdTomato/+^ mice were obtained via crossing of Rosa tdTomato (tdT) reporter mice (Jackson Laboratory, #007914) with Cx3cr1CreERT2 (Jackson Laboratory, #020940). These mice constitutively express YFP under the Cx3cr1 promoter and exposure to tamoxifen induces the expression of tdT in Cx3cr1 expressing cells. For fate mapping experiments, mice received tamoxifen via feeding with tamoxifen-containing chow (Envigo) for 10 days. The expression of YFP and tdT in Cx3Cr1 blood monocytes was confirmed by flow cytometry (BD FACSAria™ III) after tamoxifen exposure. As short-lived monocytes are constantly replenished through proliferation of bone marrow progenitors, they lose tdT expression after 21 days of washout. In contrast, resident macrophages remain tdT positive. Thus, this system can distinguish between resident cells which express both, YFP and tdT, and recruited monocyte-derived macrophages, which only express YFP. Sepsis was induced 21 days after tamoxifen exposure. Quantification of resident macrophages (Tdt-) was realized after the washout, at day 4 and day 14 after sepsis by flow cytometry and immunofluorescence.

### Immunofluorescence

Kidneys were collected, immersed in 2% formaldehyde during 6 h then in 30 % sucrose overnight at 6°C and embedded in OCT compound. Sections at 7 μm thickness (CM3050S cryostat, Leica) were stained with FITC anti-GFP antibody (ab6662, Abcam®, 1:100) and then with rabbit anti-CD146 (ab75769, Abcam, 1:100). The secondary antibody anti-rabbit coupled to DyLight™ 650 (84546, Invitrogen®, 1:100) was used.Images were acquired on a confocal microscope (LSM 700, Zeiss®) with a x20 magnification. Cells were counted blindly of group allocation per 2 by 2 tiles.

### Single-Cell Droplet Library Preparation

For scRNA-seq analysis on the 10×Genomics platform, single-cell suspensions from the kidney were prepared by pooling three animals for each group, and preparation was performed as outlined in the 10× Genomics Single Cell 3′ Reagent Kit User Guide version 2. A total of 50,000 to 100,000 live CD45+CD11b+ cells were sorted on a MACSQuant® Tyto® Cell Sorter. Samples were washed twice in PBS (Sigma), followed by centrifugation at 500 × *g* for 5 minutes at 4°C. Sample viability was assessed using trypan blue (Sigma) with an automated cell counter (Countess III), and the appropriate volume for each sample was calculated. The chip was loaded with 20,000 cells per lane. After Gel Beads-in-emulsion (GEMs) generation, samples were transferred onto a prechilled 96-well plate, heat sealed, and reverse transcription was performed using a Thermal Cycler (Bio-Rad). After reverse transcription, cDNA was recovered using the 10× Genomics Recovery Agent, and a Silane DynaBead (Thermo Fisher) cleanup was performed. Purified cDNA was amplified and cleaned using SPRIselect beads (Beckman). Samples were diluted at 4:1 (elution buffer [Qiagen]/cDNA) and an initial concentration check was performed on a Qubit fluorometer (Invitrogen) to ensure adequate cDNA concentration. The final cDNA concentration was checked on a bioanalyzer (Invitrogen). Sequencing was performed on Novaseq 6000 ILLUMINA. Using 100 cycles cartridge from ILLUMINA, with calculation based on minimum 40000 reads/cell per sample.

### Single-Cell statistical Analysis

The analysis of the single-cell data was carried out in the Data Analysis Core of Paris Brain Institute (RRID:SCR 026138). The Cell Ranger Single-cell Software suite (6.1.2) was used to process the data. Count function was used on each GEM well that was demultiplexed by mkfastq to generate gene-cell matrices. FASTQ files are aligned on the mm39 reference genome. Then, filtered_feature_bc_matrix output was loaded into Seurat package v4.3.0 to filter the datasets and identify cell types using R v4.2.2. Genes expressed in at least 5 cells and cells with at least 200 features were retained for further analysis.

All samples were merged for downstream analysis. As no batch effects were observed among the samples, no integration was performed. The gene expression matrix was normalized using the negative binomial regression method implemented in the Seurat NormalizeData() function, via selection of the top 2000 variable genes and regressed out based on the mitochondrional expression percentage. The final dataset of the kidney was composed of 17930 genes and 7424 cells. The final dataset of the heart was composed of 15265 genes and 1246 cells.

To cluster cells, we computed a Principal Components Analysis (PCA) on scaled variable genes, as determined above, using Seurat’s RunPCA function. Uniform Manifold Approximation and Projection (UMAP) was computed using Seurat’s RunUMAP function on the top 28 PCs for kidney and 19 PCs for heart. The k-nearest neighbor graph on these top PCs was computed using Seurat’s FindNeighbors function with default parameters, and the Seurat’s FindClusters function with varying resolution values using the Louvain algorithm. A value of 0.7 was chosen for the kidney clustering resolution and 0.5 for the heart resolution parameter at this stage of clustering. The FindAllMarkers function with the default parameters (min.LogFC = 0.25, min.pct = 0.25, test.use = Wilcox) was used to identify differentially expressed genes for one cluster compared to all others. Clusters were assigned preliminary identities based an automated cluster annotation by comparing with PBMC public dataset and combinations of up marker gene expression for major cell types based on literature review.

Cluster was subset without NK, B-cells, T-cells and epithelial cells. We chose a final value of 0.7 for the kidney resolution and 0.3 for the heart resolution parameter for the final clustering. Clusters were assigned preliminary identities based on expression of combinations of known marker genes for major cell types. The kidney final subset dataset was composed of 7424 genes and 17930 cells. The heart final dataset was composed of 15265 genes and 1246 cells.

Then markers were calculated for each monocyte and macrophage subtype, comparing the two conditions (sham and CLP), with the FindAllMarkers function of Seurat using default parameters. Afterwards, Gene Set Enrichment Analysis (GSEA) was run on these gene marker lists with the clusterProfiler package v4.6 using MSigDB pathways (msigdbr package v7.5.1).

## Acknowledgments

We would like to thank C. Combadières, E. Laurent-Gauthier, J. Zuber for helpful discussions, O. Thibaudeau and the UMR 1152 team for help with kidney histology. This study was conducted with the support of the Paris Brain Institute’s Data Analysis Core (https://dac.institutducerveau-icm.org/). We thank the Data Analysis Core of Sorbonne Université (RRID:SCR 026138) for preliminary analyses, in particular Riwan Brillet, Emeline Cherchame and Marie Coutelier for assistance and advice with single-cell and spatial transcriptomics data analysis.

## Author contributions

CdR and BGC designed and performed experiments with the help of VB, AB, JG, SM, MD, PLB. NT, MS, RL. Bioinformatics analyses were performed with the help of JB. CdR performed all surgeries. BGC, CdR and MD contributed to study design and data collection. CdR, AM, EG, JLS and BGC contributed to data interpretation and writing of the manuscript. All authors reviewed and approved the manuscript.

## Data and materials availability

All single-cell RNA sequencing data generated in this study have been deposited in the Gene Expression Omnibus (GEO) database and will be released on 25-06-2024. Other data will be made available upon motivated request formulated to the corresponding author.

## ONLINE SUPPLEMENTARY MATERIAL

